# Ionic Mechanisms Underlying Bistability in Spinal Motoneurons: Insights from a Computational Model

**DOI:** 10.1101/2025.06.06.658369

**Authors:** Yaroslav Molkov, Florent Krust, Russell Jeter, Tommy Stell, Mohammed A. Y. Mohammed, Frédéric Brocard, Ilya A. Rybak

## Abstract

Spinal motoneurons are the final output of spinal circuits that engage skeletal muscles to generate motor behaviors. Many motoneurons exhibit bistable behavior, alternating between a quiescent resting state and a self-sustained firing mode, classically attributed to plateau potentials driven by persistent inward currents. This intrinsic property is important for normal movement control, but can become dysregulated, causing motor function deficits, like spasticity. Here we use a conductance-based single-compartment model, together with mouse spinal slice recordings,to investigate the ionic interactions underlying motoneuron bistability. We show that synergistic interactions among high-voltage-activated L-type Ca^2+^ current (*I*_*CaL*_), calcium-induced calcium release (CICR) and the Ca^2+^-activated non-specific cation current (*I*_*CAN*_) constitute a minimal mechanistic core that produces plateau potentials and bistable firing. Within this framework, the persistent sodium current (*I*_*NaP*_) promotes plateau generation, in contrast to the Ca^2+^-dependent K^+^ current (*I*_*KCa*_) which opposes it. These results delineate ionic dependencies at the level of interactions rather than spatial localisation and provide a tractable basis for interpreting altered motoneuron excitability in disease.

**Key Points:**

- We investigated how spinal motoneurons, critical for skeletal muscle control, exhibit bistability, switching between quiet and self-sustained firing. This property stabilizes motor functions like postural control, and its dysregulation contributes to disorders such as spasticity. Using a single-compartment computational model and mouse spinal slice recordings, we explored the ionic interactions driving bistability.
- Our findings reveal that a calcium-activated cation non-specific current and calcium-induced calcium release form a core mechanism supporting the plateau depolarization essential for bistable firing. Within this framework, the persistent sodium current facilitates plateau generation, while the calcium-dependent potassium current counteracts it. Pharmacological manipulations in slices yielded results consistent with these current roles.
- Our study delineates the ionic dependencies of motoneuron bistability based on interactions, not spatial location. This offers a concise framework for interpreting excitability changes observed in normal conditions and following spinal cord injury, providing valuable insights into motor function and neurological disorders.

## Introduction

Motoneurons show a variety of nonlinear intrinsic behaviors that determine their input-output properties (Binder et al., 2020). Among them, bistability allows motoneurons to toggle between two stable states: a quiescent resting state, and a self-sustained firing state characterized by a regular pattern of action potentials generated in absence of synaptic drive. A short excitatory input can trigger a transition from the quiescent state to the active state, whereas a brief inhibitory input can induce the opposite transition (Hultborn et al., 1975; Hounsgaard et al., 1984). The underlying mechanism of the bistability involves formation of plateau potentials, long-lasting membrane depolarizations, during which the motoneuron firing is maintained (Hounsgaard, Hultborn, et al., 1988; Hounsgaard & Mintz, 1988). Persistent firing described in motor units in intact animals (Eken et al., 1989; Eken & Kiehn, 1989; Kiehn & Eken, 1997) as well as in humans (M. A. Gorassini et al., 1998; M. Gorassini et al., 1999; Collins et al., 2001) provides indirect evidence that motoneuron bistability is an important feature of motor control contributing to many behaviors. It has been suggested that bistability is involved in postural control lessening the requirement for continuous synaptic drive (Hounsgaard, Hultborn, et al., 1988; Lee & Heckman, 1998a; Heckman et al., 2009). Consistent with this, limiting motoneuron bistability was shown to impair postural control (Bos et al., 2021).

Early studies proposed that bistability primarily resulted from a persistent component of the calcium current (*I*_*CaL*_) (Hounsgaard & Mintz, 1988; Hounsgaard & Kiehn, 1989) conducted through Cav1.3 channels (Carlin et al., 2000; Simon et al., 2003; Zhang et al., 2006). However, subsequent research has shown that bistability results from a more complex interaction of several ionic currents (Bouhadfane et al., 2013), including the calcium-activated non-specific cation current (*I*_*CAN*_) through the Transient Receptor Potential Melastatin 5 (TRPM5) channels (Bos et al., 2021) and the persistent sodium current (*I*_*NaP*_) via Nav1.6 channels (Drouillas et al., 2023). The emerging picture is a causal sequence of cellular processes: the initial membrane depolarization activates *I*_*NaP*_, triggering firing and calcium influx; *I*_*CaL*_ increases intracellular calcium, which then activates calcium-induced calcium release (CICR) and recruits *I*_*CAN*_, leading to further depolarizing of the membrane. This cascade of depolarization and triggered currents leads to self-sustained firing, reinforcing the positive feedback loop underlying bistability. The contribution of these intrinsic processes to motoneuron bistability is closely tied to cell size with larger motoneurons exhibiting both higher current expression and a greater propensity for bistable behavior (Harris-Warrick et al., 2024).

Bistability is modulated by neuromodulatory and biophysical contexts. Monoamines released from supraspinal centers can promote and unmask bistability in motoneurons with its latent or partial expression (Conway et al., 1988; Hounsgaard, Hultborn, et al., 1988; Hounsgaard & Kiehn, 1989; Perrier & Delgado-Lezama, 2005). Temperature also modulates bistability: values above 30°C unmask plateau potentials by recruiting thermosensitive currents mediated by TRPM5 channels (Bos et al., 2021).

Beyond its physiological role, bistability has critical implications in pathology. Following spinal cord injury, excessive bistability in motoneurones has been linked to spasticity, a disabling condition characterized by involuntary muscle contractions (Bennett et al., 2001; Heckman et al., 2008) arising from dysregulated intrinsic ionic currents that amplify self-sustained firing and disrupt motor control (C. Brocard et al., 2016; Murray et al., 2010).

Despite significant progress, our understanding of multiple ionic mechanisms underlying bistability remains incomplete. Prior studies often examined the roles of single currents in isolation, leaving open the possible importance of their integration and interactions for motoneuron excitability. In this study, we combine computational modeling, using a conductance-based single-compartment motoneuron model, with mouse spinal slice recordings to dissect how these currents, considered separately and in combination, control the emergence, maintenance and modulation of bistability in motoneurons. Our goal is to identify a minimal mechanistic core of motoneuron bistability at the level of ionic interactions.

## Methods

### Modeling methods

#### Motoneuron model

We used a conductance-based single-compartment mathematical model of a motoneuron that includes the main spike-generating channels, fast sodium (*I*_*NaF*_) and potassium rectifier (*I*_*Kdr*_), as well as the persistent sodium (*I*_*NaP*_), slowly inactivating potassium (*I*_*Kv1*.*2*_), high-voltage activated calcium (*I*_*CaL*_), *Ca*^*2+*^-activated, non-specific cation (*I*_*CAN*_, associated with TRPM5 channels) and *Ca*^*2+*^-dependent potassium (*I*_*KCa*_, associated with SK channels) current.

The voltage dynamics are described by the current balance equation:

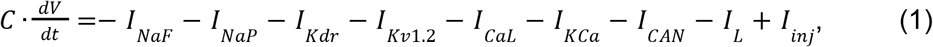

where *V* is membrane potential in mV, *t* is time in ms, *C* is membrane capacitance (*C* = 1 μ*F*·*cm* ^−2^), and the right hand side of the equation contains all transmembrane currents described as follows.

Fast sodium current (Booth et al., 1997):

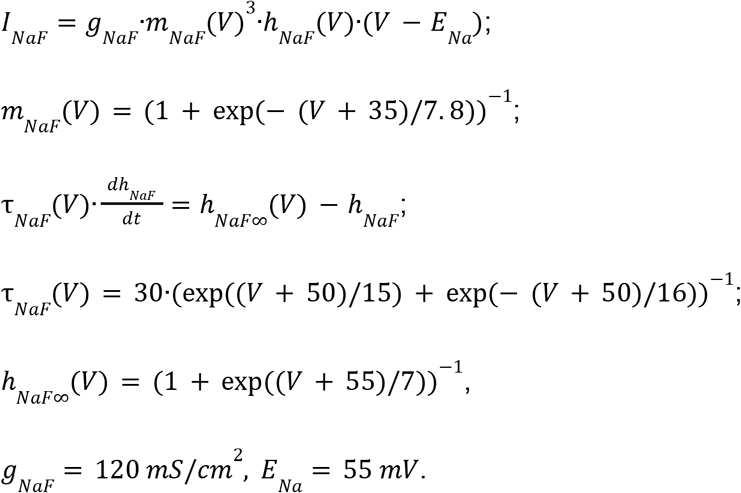

The persistent sodium current was assumed to be non-inactivating. Activation was assumed instantaneous and non-inactivating. Activation dependence on voltage was taken from (F. Brocard et al., 2013), :

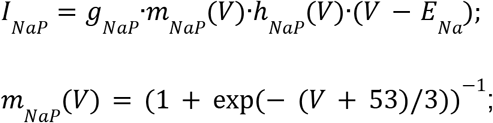

Persistent sodium current is assumed non-inactivating, i.e. *h* _*NaP*_ (*V*) = 1.

Potassium rectifier current (Booth et al., 1997):

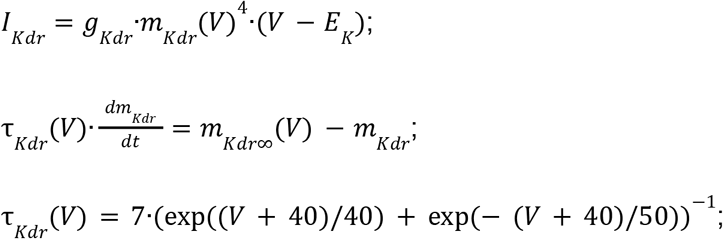

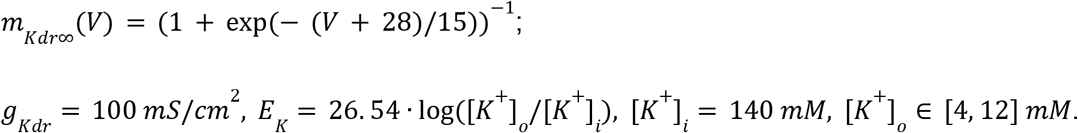

Slowly inactivating potassium current (Bos et al., 2018):

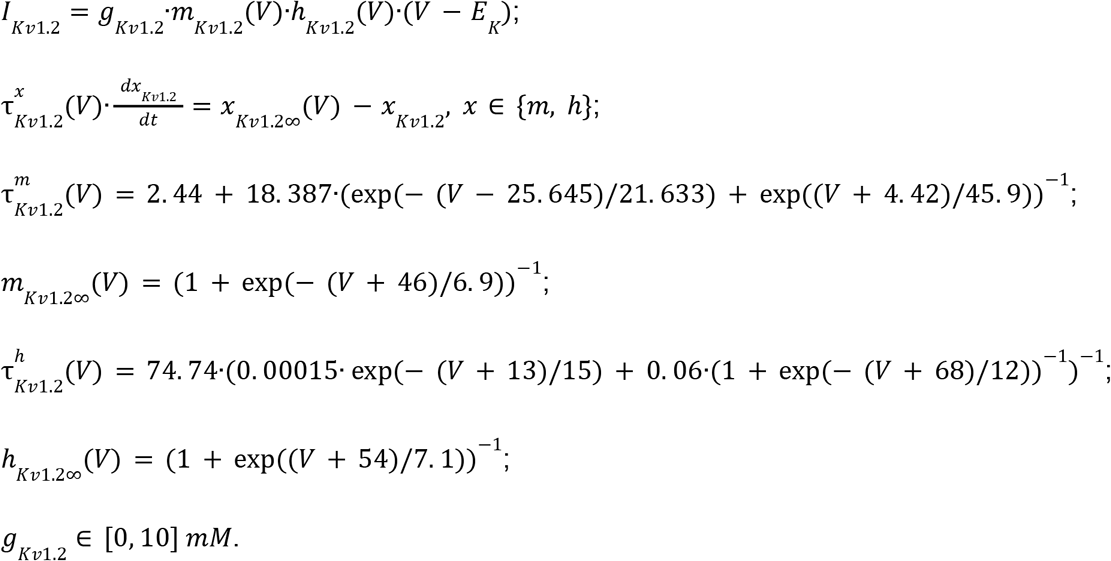

High-voltage calcium *L* (*I*_*CaL*_) current (Jasinski et al., 2013):

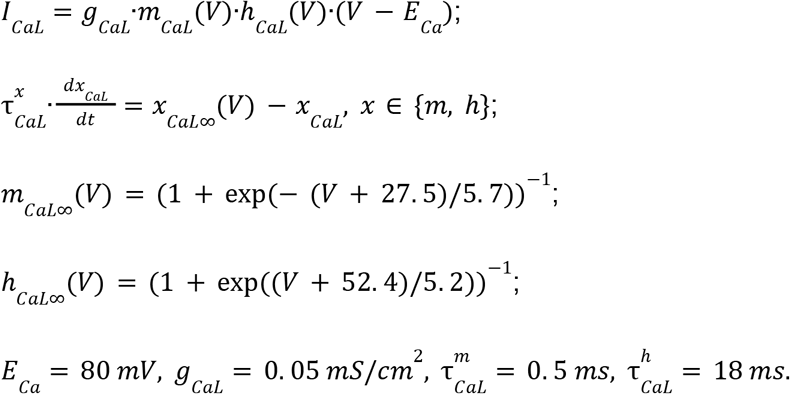

Calcium-activated nonspecific cation (*I*_*CAN*_) current (Toporikova & Butera, 2011):

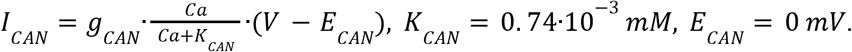

Calcium-dependent potassium current (*I*_*KCa*_) (Booth et al., 1997). This current was modified to be instantaneous:

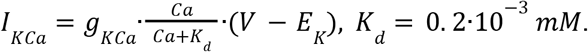

Leak current:

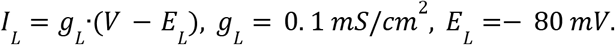

The two currents described above (*I*_*CAN*_ and *I*_*KCa*_) depend on the intracellular *Ca*^*2+*^ concentration [Ca^2+^]_i_ in mM (denoted by *Ca* in the equations). The *Ca*^*2+*^ concentration increases directly from the influx of calcium ions through calcium channels (captured by *I*_*CaL*_ in the mathematical model) and indirectly from the release of calcium ions from intracellular stores via a calcium-induced calcium release (CICR) mechanism. Additionally, they are pumped out by the Ca-ATP pumps.

The dynamics of intracellular calcium concentration (*Ca*) in our model are described by the differential equation:

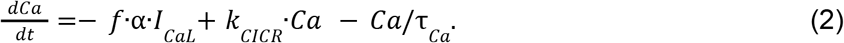

On the right hand side of this equation, the first term represents calcium influx through high-threshold, voltage-gated calcium channels (*I*_*CaL*_) which open during action potentials. Here *f* = 0. 01 defines the ratio of entered Ca^2+^ ions remaining unbound; the coefficient *α* = (2·*F*·*δ*)^−1^ converts inward *I*_*CaL*_ current to *Ca*^*2+*^ concentration rate of change; here *F* is

Faraday’s constant (*F* = 9. 648 ·10^4^ *C*/*mol*) and *δ* is the thickness (0. 1 μ*m*) of the shell adjacent to the membrane. Based on these parameters, *α* = 5·10 ^−4^*mM*·*cm*^2^ ·*ms*^−1^ ·μ*A*^−1^.

The second term describes an increase of cytoplasmic *Ca*^*2+*^ concentration through the CICR mechanism, where the rate of calcium release from internal stores is proportional to the intracellular *Ca*^*2+*^ concentration (defined by *k*_*CICR*_). The third term describes the action of calcium pumps (both plasma membrane and SERCA) which rapidly remove calcium from the cytoplasm, with a time constant *τ* _*Ca*_ = 10 *ms*.

#### Qualitative analysis

To investigate the bistable behavior of spinal motoneurons, we implemented a current ramp simulation protocol designed to probe the transitions between quiescent and self-sustained firing states. This approach leverages a linearly varying injected current (*I*_*inj*_) to systematically explore the system’s response across a range of input intensities, making it an effective tool for detecting hysteresis and state-dependent dynamics in neuronal models.

The protocol was designed as follows: the injected current was initially set to zero and then increased linearly to a predetermined maximum value (ascending phase) over a specified duration. Subsequently, the current was decreased linearly back to zero (descending phase) at the same rate. This bidirectional ramp allowed us to identify two key transition points: the current threshold at which the system shifts from silence to spiking during the ascending phase (*I*_*up*_) and the lower threshold at which spiking ceases during the descending phase (*I*_*down*_). Bistability is indicated when *I*_*down*_ is less than *I*_*up*_, revealing a range of current values where the system can stably maintain either state, depending on its prior condition. This hysteresis reflects the nonlinear properties of the model and its history-dependent behavior. To ensure the reliability of these thresholds, the ramp time was varied systematically, extending the duration of the ascending and descending phases until *I*_*up*_ and *I*_*down*_ stabilized, confirming that the observed transitions were not influenced by transient dynamics. Typically, ramp durations spanned several hundred milliseconds, with longer durations tested to exclude such effects.

To further validate the protocol’s ability to detect bistability, we employed a complementary current step simulation inspired by experimental approaches. Starting from a baseline of *I*_*inj*_ = 0, the current was stepped to an intermediate value within the suspected bistable range (*I*_*down*_ < *I*_*inj*_

< *I*_*up*_), then increased to a level above *I*_*up*_, and subsequently returned to the intermediate value before returning to zero. This sequence demonstrated that, at the intermediate current level, the system’s state—silent or spiking—depended on its prior activation history, reinforcing the findings of the ramp protocol.

The current ramp simulations also supported parametric analyses by varying key model parameters and plotting the resulting bifurcation diagrams. These diagrams mapped the system’s equilibrium points and oscillatory regimes as functions of *I*_*inj*_, providing a visual representation of the bistable region. The protocol’s design, with its carefully adjusted ramp duration, ensured that the gradual variation of input current effectively captured the boundaries of this region, offering a robust technical framework for studying the conditions under which bistability emerges and persists in the model.

#### Simulations

Simulations were performed using custom-written C++ and Julia software. Integration was performed by the Dormand-Prince 5(4) method using (*Boost C++ Libraries*, ver. 1.86). Source code written in C++ and Julia for the model and examples of the ramping protocols can be found in the Github repository associated with this manuscript (Jeter & Molkov, 2025).

### Experimental methods

#### Experimental model

Mice (C57/Bl6 background) were housed under a 12h light/dark cycle with *ad libitum* access to water and food. Room temperature was kept between 21-24°C and between 40-60% relative humidity. All animal care and use were conformed to the French regulations (Décret 2010-118) and approved by the local ethics committee (Comité d’Ethique en Neurosciences INT-Marseille, CE71 Nb A1301404, authorization Nb 2018110819197361).

#### In vitro preparations

For the slice preparation, the lumbar spinal cord was isolated in ice-cold (+4°C) artificial CSF (aCSF) solution composed of the following (in mM): 252 sucrose, 3 KCl, 1.25 KH2PO4, 4 MgSO4, 0.2 CaCl2, 26 NaHCO3, 25 D-glucose, pH 7.4. The lumbar spinal cord was then introduced into a 1% agar solution, quickly cooled, mounted in a vibrating microtome (Leica, VT1000S) and sliced (325 µm) through the L4–5 lumbar segments. Slices were immediately transferred into the holding chamber filled with bubbled (95% O2 and 5% CO2) aCSF solution composed of (in mM): 120 NaCl, 3 KCl, 1.25 NaH2PO4, 1.3 MgSO4, 1.2 CaCl2, 25 NaHCO3, 20 D-glucose, pH 7.4, 30-32°C. After a 30-60 min resting period, individual slices were transferred to a recording chamber continuously perfused with aCSF heated to 32-34°C.

#### In vitro recordings

Whole-cell patch-clamp recordings were performed using a Multiclamp 700B amplifier (Molecular Devices) from L4-L5 motoneurons with the largest soma (>400µm2) located in the lateral ventral horn. Patch electrodes (2-4 MΩ) were pulled from borosilicate glass capillaries (1.5 mm OD, 1.12 mm ID; World Precision Instruments) on a Sutter P-97 puller (Sutter Instruments Company) and filled with an intracellular solution (in mM): 140 K+-gluconate, 5 NaCl, 2 MgCl2, 10 HEPES, 0.5 EGTA, 2 ATP, 0.4 GTP, pH 7.3. Pipette and neuronal capacitive currents were canceled and, after breakthrough, the series resistance was compensated and monitored. Recordings were digitized on-line and filtered at 20 kHz through a Digidata 1550B interface using Clampex 10.7 software (Molecular Devices). All experiments were designed to gather data within a stable period (i.e., at least 2 min after establishing whole-cell access).

#### Data quantification

Electrophysiological data analyses were analyzed off-line with Clampfit 10.7 software (Molecular Devices). For intracellular recordings, several basic criteria were set to ensure optimum quality of intracellular recordings. Only cells exhibiting a stable resting membrane potential, access resistance (no > 20% variation) and an action potential amplitude larger than 40 mV under normal aCSF were considered. Passive membrane properties of cells were measured by determining from the holding potential the largest voltage deflections induced by small current pulses that avoided activation of voltage-sensitive currents. We determined input resistance by the slope of linear fits to voltage responses evoked by small positive and negative current injections. The peak amplitude of the slow afterdepolarization (slow ADP or sADP) was defined as the difference between the holding potential and the peak voltage deflection after the burst of spikes. The sADP area was measured between the end of the stimulus pulse and the onset of the hyperpolarizing pulse (delta= 7.5 s). In addition, the duration of the sADP was quantified as the time interval at half-maximal amplitude. If necessary, using bias currents, the pre-pulse membrane potential was maintained at the holding potential fixed in the control condition. Bistable properties were investigated with a 2 s depolarizing current pulses of varying amplitudes (0.8 - 2 nA). To quantify the ability of a motoneuron to be bistable, we gradually increased the holding current in 25 pA steps over a 2-second period before delivering the depolarizing pulse. This process continued until the neuron reached its spiking threshold. The cell was considered as bistable when (1) the pre-stimulus membrane potential stays relatively hyperpolarized below the spiking threshold (downstate), (2) the post-stimulus membrane potential stays depolarized above the spike threshold (upstate), and (3) the membrane potential switches to downstate after a brief hyperpolarizing pulse. To quantify the extent of bistability, we measured both the voltage (*V*) range between the most hyperpolarized holding potential (*Vh min*) and the most depolarized holding potential (*Vh max)* at which the motoneuron can exhibit a self-sustained spiking, and the corresponding range of injected currents (*I*) over which bistable behavior was observed.

#### Statistics

When two conditions (control vs drugs) were compared, we used the Wilcoxon matched pairs test. For all statistical analyses, the data met the assumptions of the test and the variance between the statistically compared groups was similar. The level of significance was set at p < 0.05. Statistical analyses were performed using Graphpad Prism 7 software.

## Results

### Calcium dynamics and calcium-dependent currents

#### The role of calcium-induced calcium release (CICR) mechanism

A brief excitatory current pulse into a motoneuron triggers a train of action potentials, causing a substantial increase in intracellular calcium levels ([Ca^2+^]_i_). This increase is driven by influx via voltage-gated calcium channels, further amplified by calcium-induced calcium release (CICR) from internal stores. The resulting [Ca^2+^]_i_ accumulation activates two Ca^2+^-dependent currents with opposing effects: the depolarizing calcium-activated nonspecific cation current (*I*_*CAN*_) and the hyperpolarizing calcium-dependent potassium current (*I*_*KCa*_).

As described in Methods, the [Ca^2+^]_i_ dynamics in our model are governed by the differential equation (2). The calcium clearance pump operates with a time constant of *τ* _*Ca*_ = 10 ms, which, in the absence of CICR, would clear all calcium introduced by a spike before the next spike. The term *k*_*CICR*_ ·*Ca* in equation (2) captures the CICR mechanism, where the rate of calcium release from internal stores is directly proportional to the current calcium concentration *Ca* with coefficient *k*_*CICR*_.

For simplicity, the intracellular calcium dynamics described by Eq. (2) can be expressed as:

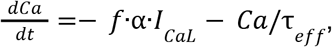

where *τ*_*eff*_ represents the effective time constant

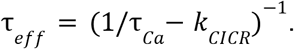

The gain *k*_*CICR*_ must not exceed 1/*τ*_*Ca*_ to prevent negative *τ*_*eff*_, which would lead to an infinite increase in calcium. In our model, *k*_*CICR*_ is set at 0.096 ms^-1^, yielding an effective time constant *τ* _*eff*_ = 250 ms. This value was chosen phenomenologically to match typical slow afterdepolarization decay rates observed in our *in vitro* data and prior studies.

This prolonged *τ*_*eff*_ (compared to *τ*_*Ca*_) reflects how the CICR mechanism substantially slows calcium clearance, resulting in [Ca^2+^]_i_ build-up during repetitive firing. As shown in Figure 1, calcium levels sharply increase in response to a rectangular current pulse, strongly activating *I*_*CAN*_ and *I*_*KCa*_. Without CICR (*k* _*CICR*_ = 0), pumps clear calcium rapidly between spikes, preventing significant accumulation during repetitive firing. Consequently, in the absence of sustained elevations in intracellular *Ca*^*2+*^, neither *I*_*CAN*_ nor *I*_*KCa*_ currents are activated to a significant degree.

**Figure 1.**
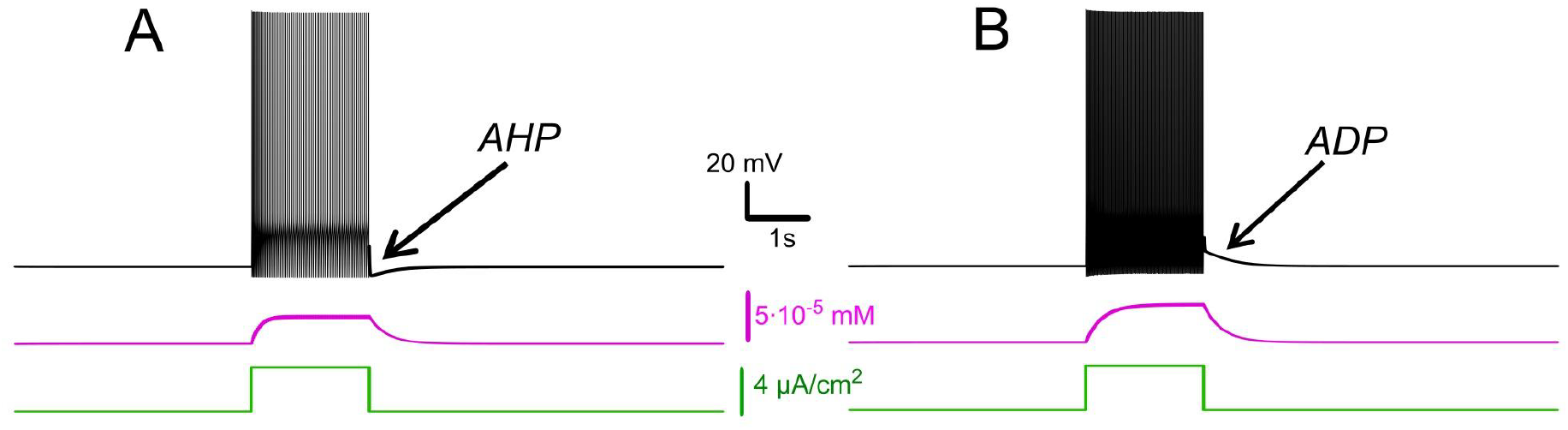
Effects of *I*_*KCa*_ vs. effects of *I*_*CAN*_. Membrane potential (black) and intracellular *Ca*^*2+*^ (magenta) in response to step current (green). Intracellular *Ca*^*2+*^ concentration increases during spiking activity and then slowly adapts. **A.**When *I*_*KCa*_ is stronger than *I*_*CAN*_ (*g*_*KCa*_=0.5 mS, *g*_*CAN*_=0 mS), this leads to after-hyperpolarization (AHP). **B**. When *I*_*CAN*_ is stronger than *I*_*KCa*_ (*g*_*KCa*_=0.5 mS, *g*_*CAN*_=0.7 mS), this leads to after-depolarization (ADP) immediately following spiking activity.

#### Interplay between *I*_*CAN*_ and *I*_*KCa*_: afterdepolarization versus afterhyperpolarization

Intracellular recordings have shown that motoneurons demonstrate two post-stimulus behaviours: a slow afterdepolarization (sADP) after brief excitatory inputs, typical for a bistable motoneuron type, and an afterhyperpolarization (AHP) observed in a non-bistable motoneuron type (Harris-Warrick et al., 2024). Our model captures both behaviors and links them to the relative contributions of *I*_*CAN*_ and *I*_*KCa*_. We also show that these currents have opposite effects on motoneuron bistability. When *I*_*KCa*_ predominates, *Ca*^*2+*^ -dependent potassium efflux produces hyperpolarization of the membrane and a decrease in excitability. This effect promotes spike-frequency adaptation and a prominent post-stimulus AHP supported by sustained [Ca^2+^]_i_ elevation (Fig. 1A).

Conversely, when *I*_*CAN*_ predominates, the elevation of [Ca^2+^]_i_ activates this current, leading to a depolarizing influx of sodium, that in turn increases the firing frequency during the pulse and yields an sADP afterward (Fig. 1B). As [Ca^2+^]_i_ declines, the membrane potential slowly relaxes back toward its resting values. These model behaviours are consistent with recordings and provide a mechanistic interpretation: *I*_*CAN*_ biases toward sADP and plateau maintenance, whereas *I*_*KCa*_ biases toward AHP and termination of spiking.

#### Mechanisms of *I*_*CAN*_-based bistability and its modulation *I*_*CAN*_-based bistability

Our model predicts that the expression of *I*_*CAN*_ provides a robust route to bistability. The process begins with the opening of voltage-gated calcium channels during each action potential. While this influx alone is insufficient to fully activate *I*_*CAN*_, it triggers CICR, which amplifies [Ca^2+^]_i_ (see below).

The elevated [Ca^2+^]_i_ activates *I*_*CAN*_ establishing a positive feedback loop: *I*_*CAN*_ sustains depolarization, promoting continuous firing; spikes further elevate [Ca^2+^]_i_ via Ca^2+^ entry and CICR, and thus reinforce *I*_*CAN*_ activation. This self-perpetuating mechanism allows the motoneuron to remain in a high-activity state (persistent spiking) even after the initial stimulus is removed, creating bistability: the neuron can operate in either a quiescent state or an active spiking state.

To validate this mechanism, we replicated the experimental ramp current protocol in the model (see Methods). At *g* = 0.5 *mS*/*cm* ^2^ the voltage–current relation during a linear *I*_*inj*_ ramp (Fig. 2C) reveals distinct up and down thresholds. For *I*_*inj*_ < *I*_*up*_ = 1.7 μ*A*/*cm*^2^, the system displays two attracting regimes separated by an unstable equilibrium (saddle): a stable hyperpolarized state corresponding to the resting state (silent) and a stable spiking state (limit cycle) (Fig. 2C). As *I*_*inj*_ exceeds *I*_*up*_ = 1.7 μ*A*/*cm* ^2^, the low potential stable branch of the *V*-nullcline merges with the saddle and vanishes via a fold bifurcation, and the system transitions to a stable limit cycle representing a repetitive spiking regime. During the descending phase of the ramp, spiking persists until *I*_inj_ decreases to *I*_*down*_ = 1.1 μ*A*/*cm* ^2^, which is less than *I*_*up*_ (Figs. 2C, D), yielding a hysteresis interval (*I*_*down*_< *I*_*inj*_ < *I*_*up*_), indicative of bistability.

**Figure 2.**
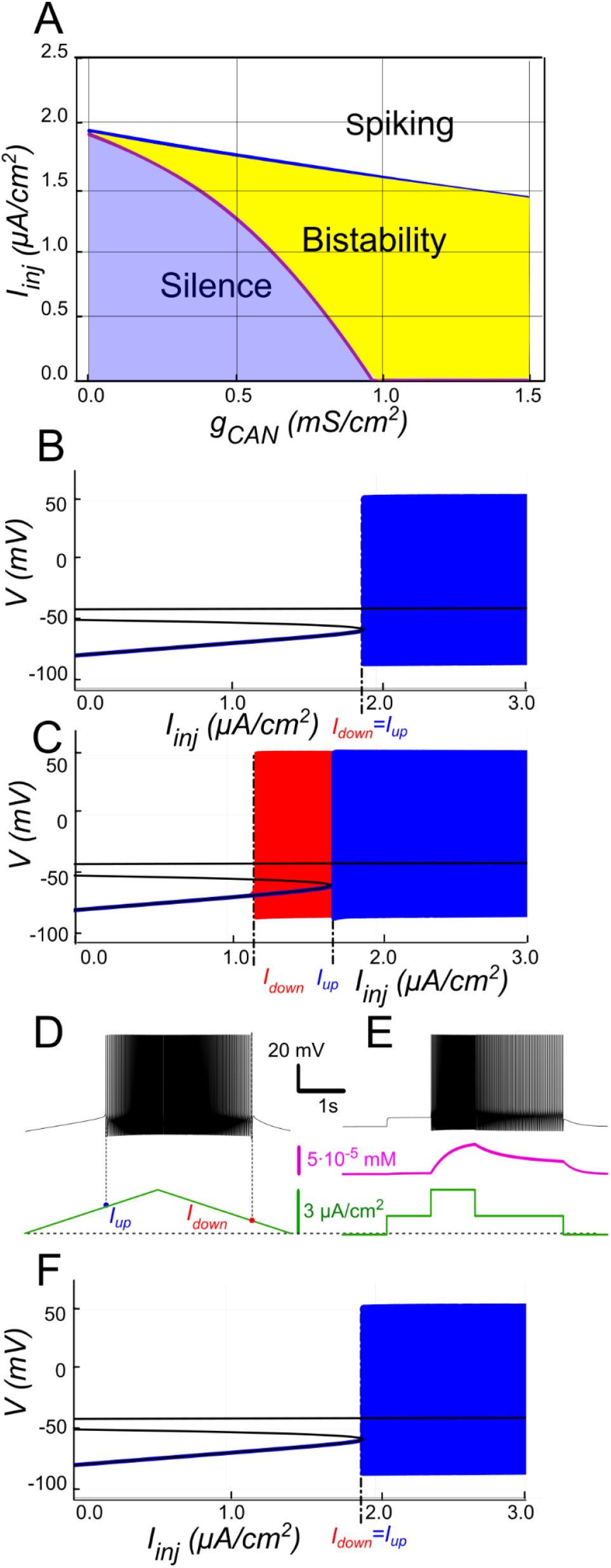
*I*_*CAN*_ -induced bistability and its disappearance after CIRC blockade. **A:** Parameter plane (*g*_*CAN*_, *I*_*inj*_) partitioned into regions of different behaviors. The upper and lower boundaries of the bistability region represent the dependence of *I*_*up*_ (blue line) and *I*_*down*_ (red line) on *g*_*CAN*_, respectively. With no *I*_*CAN*_ (*g*_*CAN*_ = 0) the transitions from silence to spiking and back occur at the same *I*_*inj*_ values signifying no bistability. As *g*_*CAN*_ increases, *I*_*down*_ becomes smaller than *I*_*up*_, with the bistability range progressively expanding. **B, C**: **Bifurcation diagrams showing different behaviors of the system** at *g*_*CAN*_ = 0 and *g*_*CAN*_ = 0.5 *mS/cm*^*2*^ across a range of the injected current values, *I*_*inj*_. Black line depicts the system’s equilibrium states. When *I*_*inj*_ < *I*_*up*_ there is a stable hyperpolarized state (stable node). If *I*_*inj*_ is increased over *I*_*up*_, the system transitions to spiking, covering the voltage range shown in red and blue. This stable spiking regime exists in the range *I*_*inj*_ > *I*_*down*_. When *I*_*inj*_ is reduced below *I*_*down*_, the limit cycle representing spiking disappears, and the system transitions to the low voltage stable fixed point. Between these bifurcation points the hyperpolarized state (silence) coexists with the stable limit cycle (spiking) shown in red. **D, E: Bistability revealed with ramp and step protocols. D**: *I*_*inj*_ was linearly increased from 0 to 3 *μA/cm*^*2*^ and back (green). Note that spiking started at higher current than the transition back to silence (*I*_*up*_ > *I*_*down*_). **E**: *I*_*inj*_ was held piecewise constant at 0 first, then increased to 1.5 *μA/cm*^*2*^, then to 3, then reduced back to 1.5, and, finally, to 0 (see the black trace at the bottom). At *I*_*inj*_ = 1.5 *μA/cm*^*2*^ the system exhibits spiking or silence depending on whether it was active or not during the previous stage. Note the difference in the intracellular calcium concentration levels (green trace). **F.**Same representation as in C but with intracellular calcium release blocked (*k*_*CIRC*_ = 0). Note lack of bistability.

To further probe bistability, we implemented a step protocol inspired by experimental methods. Starting at *I*_*inj*_ = 0 in a silent state, we applied an intermediate pulse within the bistable range (*I*_*down*_ < *I*_*inj*_ < *I*_*up*_), then increased it above *I*_*up*_ to induce firing, before returning to the intermediate current (Fig. 2E). This protocol showed that, at the same intermediate current value, the motoneuron could either continue firing or remain silent, depending on whether it was previously activated.

The 2-parameter bifurcation diagram (Fig. 2A) shows how *I*_*CAN*_ controls the extent of the bistability region. At *g*_*CAN*_ = 0, *I*_*down*_ and *I*_*up*_ coincide (no hysteresis; Fig. 2B). Once *g*_*CAN*_ becomes large enough, the interval (*I*_*down*_< *I*_*inj*_ < *I*_*up*_) opens and bistability emerges. This is evident as the current required to trigger spiking during the ramp-up exceeds the current at which spiking stops during ramp-down (Fig. 2C). This hysteresis widens with further increases in *g*_*CAN*_ (Fig. 2A), reflecting the strengthening of the *I*_*CAN*_-mediated positive feedback loop, which supports the self-sustained spiking state. These findings identify *I*_*CAN*_ as a key determinant of the bistable behavior.

#### The role of CICR

The model demonstrates that Ca^2+^ influx through voltage-gated calcium channels alone does not raise [Ca^2+^]_i_ enough to activate *I*_*CAN*_. To illustrate the contribution of CICR we set *k*_*CICR*_ to 0, which reduces the effective decay time constant *τ*_*eff*_ to *τ*_*Ca*_ = 10 *ms*. As shown in Fig. 2F, the transient [Ca^2+^]_i_ increase is then sharply curtailed, *I*_*CAN*_ remains weak, and the *I*_*CAN*_-based bistability collapses (ramp hysteresis vanishes). In other words, CICR effectively slows Ca^2+^ clearance (*τ*_*eff*_ in the hundreds of ms) and is necessary in this framework to maintain the Ca^2+^-dependent depolarizing drive provided by *I*_*CAN*_.

#### The role of *I*_*KCa*_ in modulating *I*_*CAN*_-based bistability

Our model indicates that *I*_*KCa*_ has a negative effect on *I*_*CAN*_ -based bistability. As [Ca^2+^]_i_ increases, SK-type K^+^ channels (mediating *I*_*KCa*_) open, allowing intracellular K? ions to exit the cell. This outward current hyperpolarizes the membrane, decreasing excitability, and preventing the self-sustaining depolarization provided by *I*_*CAN*_. Bifurcation analyses (Fig. 3) summarize this interplay. At low *g*_*CAN*_, a regime where *I*_*KCa*_ dominates the Ca^2+^-dependent response, the neuron transitions between spiking and silence at the same current threshold during ascending and descending current ramps, with this common threshold only weakly influenced by *g*_*CAN*_ (Fig. 3A). Increasing *g*_*CAN*_ lowers the spiking onset threshold and, once *g*_*CAN*_ reaches approximately 1 *mS/cm*^*2*^ (Fig. 3A), a hysteresis interval (*I*_*down*_< *I*_*inj*_ < *I*_*up*_) opens, indicating the emerging bistability. This is evident as the current required to trigger spiking during the ramp-up exceeds the current at which spiking stops during ramp-down, with the bistability range widening as *g*_*CAN*_ increases further. Additionally, the specific *g*_*CAN*_ value at which this bifurcation occurs depends linearly on *g*_*KCa*_ (Fig. 3B), suggesting that the bistability arises when *I*_*CAN*_ begins to dominate over *I*_*KCa*_.

**Figure 3.**
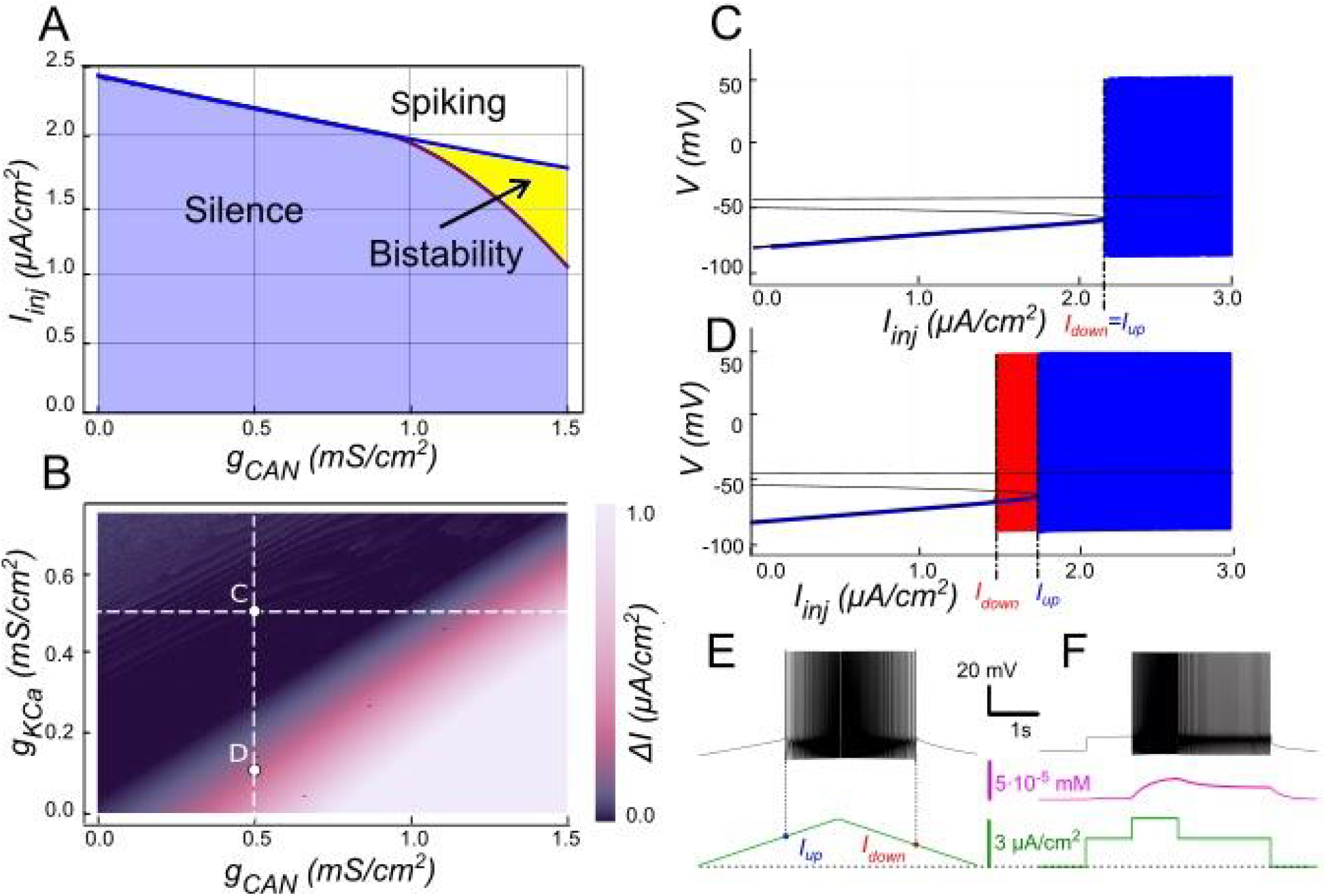
Modulation of *I*_*CAN*_-dependent bistability by *I*_*KCa*_. **A.**Bifurcation diagram similar to Fig. 2A, constructed for *g*_*KCa*_ = 0.5 *mS/cm*^*2*^. Note, that unlike in Fig. 2, bistability emerges once *g*_*CAN*_ exceeds 1 *mS/cm*^*2*^. **B**. Bistability range *ΔI*_*inj*_ depending on *g*_*CAN*_ and *g*_*KCa*_ conductances. Bistability range (defined as *I*_*up*_ - *I*_*down*_) is color coded. Bistability exists in the lower right part of the diagram. Note near linear dependence of *g*_*CAN*_ bistability threshold on *g*_*KCa*_. White dashed line shows the value of *g*_*KCa*_ used to construct the bifurcation diagram in panel A. **C, D**. Bifurcation diagrams showing possible behaviors of the system at the parameter values labelled correspondingly in panel B. **C**. At higher *g*_*KCa*_ value (*g*_*KCa*_ = 0.5) transitions from quiescence to spiking and back occur at the same injected current value (*I*_*up*_ = *I*_*down*_), indicating no bistability. **D**. When *g*_*KCa*_ is lowered to 0.1, the transition from spiking to quiescence occurs at a lower injected current than the transition from quiescence to spiking (*I*_*down*_ < *I*_*up*_), so spiking shown in red coexists with the silent regime. **E**. Ramp (left) and step (right) current injection protocols, illustrating bistability revealed in D. The intermediate current step is between *I*_*down*_ and *I*_*up*_. The system’s state depends on whether it was active or not at the previous step, exhibiting bistable behavior.

Equivalently, reducing *g*_*KCa*_ at fixed *g*_*CAN*_ unmasks bistability (Figs. 3C-F).

We tested these predictions by performing patch-clamp recordings of lumbar motoneurons, focusing on the effects of apamin, a selective blocker of *I*_*KCa*_. We measured the parameters of the sADP induced by a brief depolarization of the motoneurons. Application of apamin significantly increased the amplitude, duration, and area of the sADP, indicating a stronger and more prolonged depolarizing response when *I*_*KCa*_ is reduced (Fig. 4A-D). In addition, apamin also enhanced the capacity of motoneurons for bistable behavior. Specifically, motoneurons were able to express plateau potentials from more hyperpolarized holding potentials (Fig. 4E-F), reflected by a significant shift in Vh min from -62.6 mV to -64.8 mV (p < 0.05; Wilcoxon matched pairs test), and by increases in both the voltage range (*ΔV*) and the current range (*ΔI)* over which bistability was observed (Fig. 4G-H). Together, these results support the prediction of the model that reducing *I*_*KCa*_ expands the bistable regime by reducing the after-hyperpolarizing influence opposing *I*_*CAN*_-driven depolarization (Figs. 1 and 3). These results underscore the antagonistic interplay between *I*_*CAN*_ and *I*_*KCa*_ in shaping motoneuron excitability and provide a mechanistic basis for bistable behaviors.

**Figure 4.**
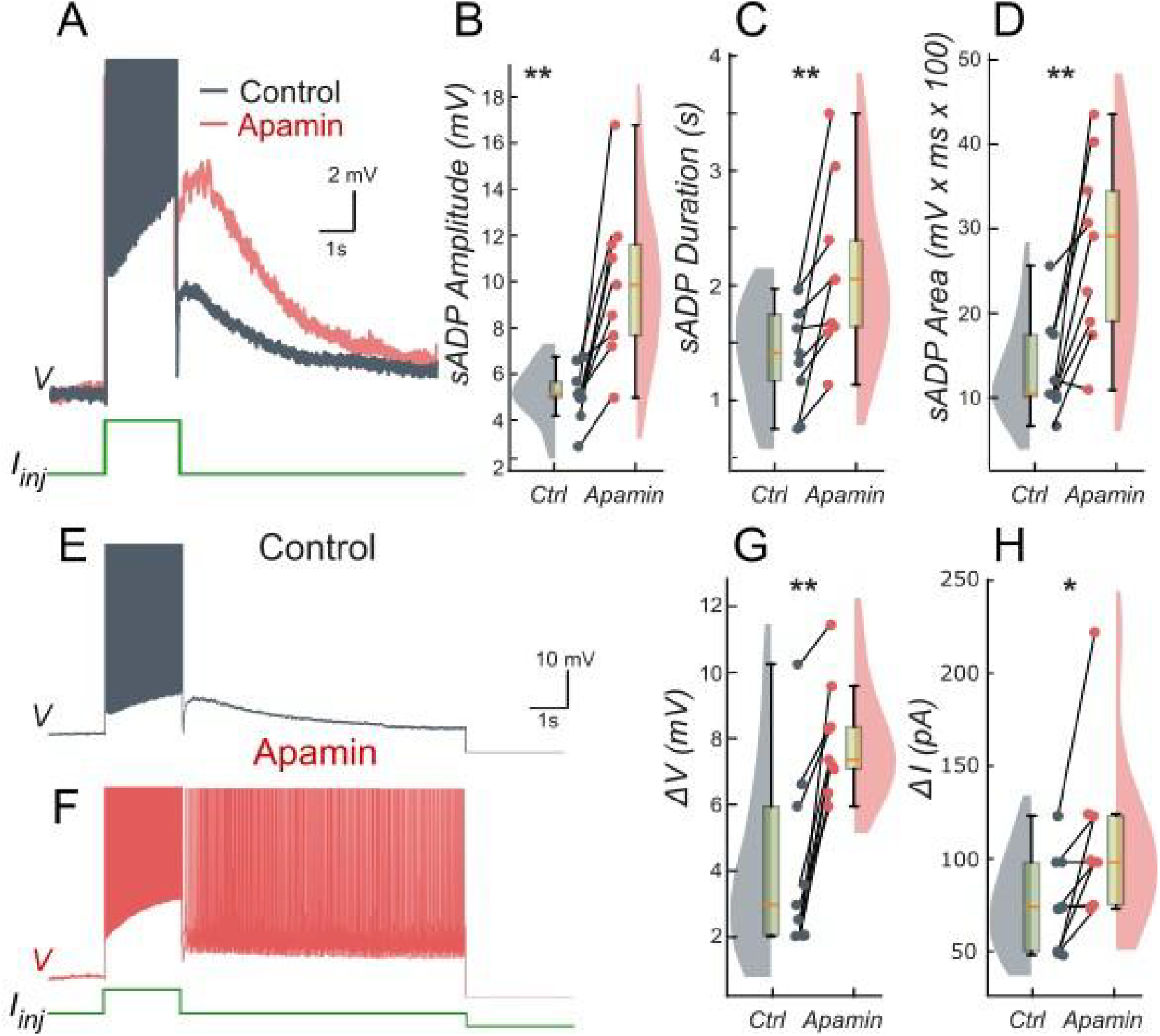
Ca^2+^-activated K+ current (*I*_*KCa*_) limits the slow afterdepolarization and membrane bistability in lumbar motoneurons. **A**: Superimposed voltage traces recorded in the same motoneuron during a brief (2 s) depolarizing current step (bottom) under control conditions (black) and after bath application of apamin (red). **B-D**: Quantification of the sADP amplitude (**B**), duration(**C**) and area (**D**). **E, F**: Voltage traces in response to a 2-s depolarizing pulse before (**E**) and after apamin (**F**). **G, H:** Quantification of bistability through ΔV and ΔI. ΔV and ΔI represent the range of holding potentials and holding currents, respectively, over which self-sustained firing can be observed. Each is defined as the difference between the most depolarized and the most hyperpolarized value (potential for ΔV, current for ΔI) at which self-sustained firing is triggered or maintained (see Methods). Paired data from individual motoneurons (n = 9) are linked and overlaid on violin and box-and-whisker plots. *P < 0.05, **P < 0.01, two-tailed Wilcoxon signed-rank test.

#### Modulation of *I*_CAN_-based bistability by extracellular potassium concentration

Because *I*_*KCa*_ depends on the *K*^*+*^ driving force, *I*_*CAN*_ -dependent bistability should be sensitive to the potassium reversal potential (*E*_*K*_), which is set by the Nernst equation. This equation provides the potassium ion equilibrium potential based on the ratio of intracellular [K^+^]_i_ to extracellular [K^+^]_o_ potassium concentration (see Methods). Increasing extracellular potassium [K^+^]_o_ depolarizes *E*_*K*_ thereby reducing the outward driving force through *I*_*KCa*_. This weakens the hyperpolarizing influence of *I*_*KCa*_ that counteracts the *I*_*CAN*_-mediated depolarizing feedback (see above).

We explored this interaction by varying [K^+^]_o_ and *g*_*CAN*_ in the model while holding *g*_*KCa*_ at 0.5 mS/cm^2^. The resulting two-parameter map (Fig. 5B) shows the bistable range (color-coded width *ΔI = I*_*up*_ *– I*_*down*_, black indicating no bistability) as a function of *g*_*CAN*_ and [K^+^]_o_. At low [K^+^]_o_ (e.g., physiologically normal levels of 4 mM), *E*_*K*_ is strongly negative and *I*_*KCa*_ efficiently counteracts *I*_*CAN*_, so relatively large *g*_*CAN*_ (e.g., ∼1 mS/cm^2^, Fig. 3A) is required for bistability. As [K^+^]_o_ increases, *E*_*K*_ depolarizes, weakening *I*_*KCa*_ and the bistable interval opens at progressively lower *g*_*CAN*_ (Fig. 5A). For instance, at *g*_*CAN*_ = 0.9 mS/cm^2^ the model is not bistable at [K^+^]_o_ = 4 mM (Fig. 5C), but becomes bistable at [K^+^]_o_ = 8 mM (Figs. 5D, E), with the hysteresis width further expanding as [K^+^]_o_ rises (Fig. 5B). As elsewhere, we corroborated hysteresis with a step protocol showing different coexisting stable regimes at identical inputs (Fig. 5F).

**Figure 5.**
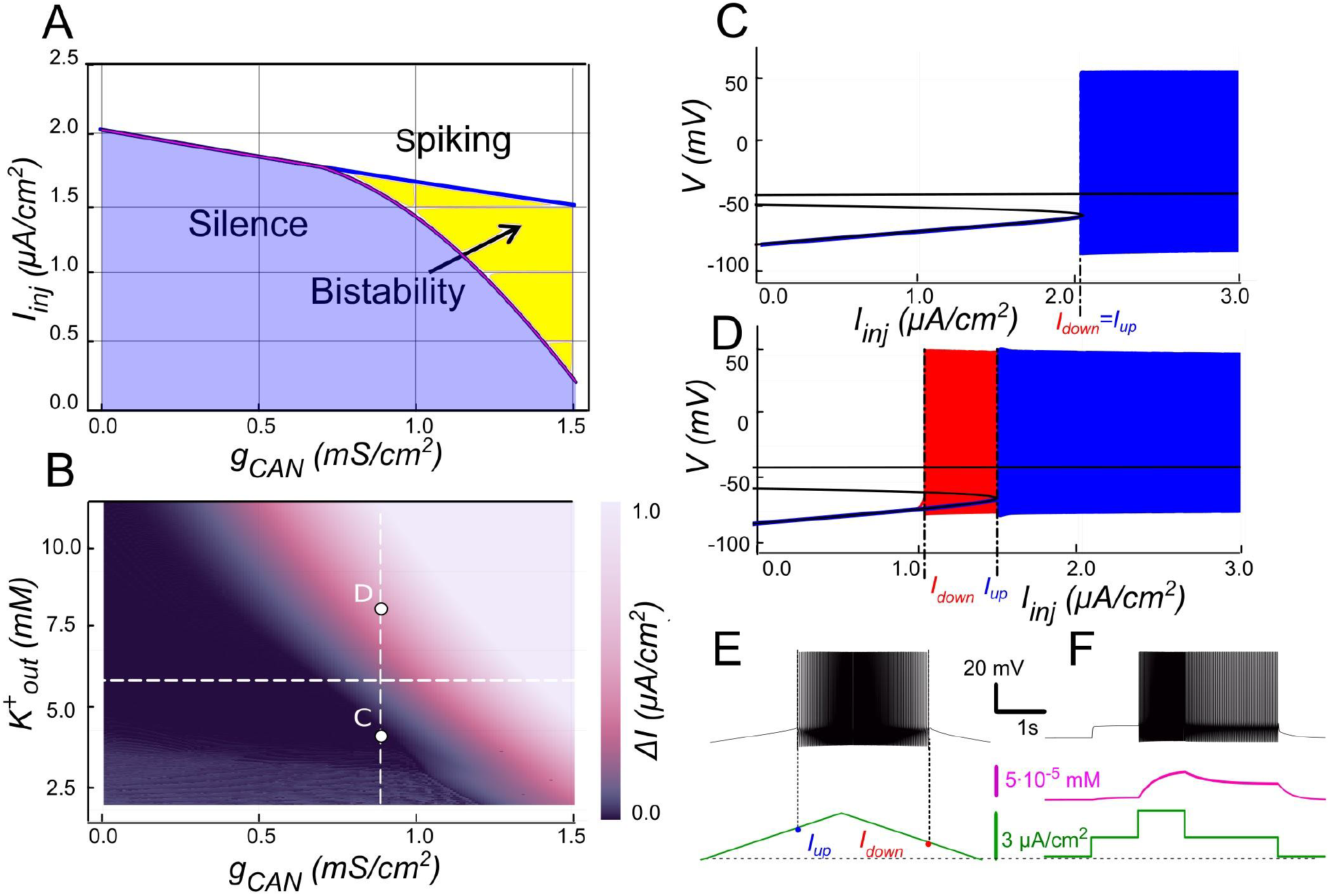
Modulation of *I*_*CAN*_-dependent bistability by extracellular potassium concentration ([K^+^]_o_). **A.**Bifurcation diagram similar to Fig. 3A, constructed for *g*_*KCa*_ = 0.5 *mS/cm*^*2*^, but at elevated [K^+^]_o_ = 6mM instead of physiologically normal 4mM. Note, that compared to Fig. 3A, bistability emerges at lower *g*_*CAN*_. **B**. Color-coded bistability range (*I*_*up*_ - *I*_*down*_) depending on *g*_*CAN*_ and [K^+^]_o_. Black area corresponds to no bistability. White dashed line shows the [K^+^]_o_ value used in panel A. *g*_*CAN*_ bifurcation value reduces as [K^+^]_o_ increases, therefore an increase in [K^+^]_o_ can lead to bistability emergence, as shown in **C** and **D**. If *g*_*CAN*_ = 0.9 *mS/cm*^*2*^, at [K^+^]_o_ = 4 mM (physiologically normal value) no bistability exists (**B**), but if [K^+^]_o_ is raised to 8 mM, bistability emerges (**C**), as illustrated by ramp (**E**) and step (**F**) current protocols.

This modulation is physiologically relevant: during sustained activity or in certain pathological conditions, elevation of [K^+^]_o_ can occur, which, by decreasing the effectiveness of *I*_*KCa*_, would favour expression of the *I*_*CAN*_-driven positive feedback and broaden the bistable operating range. More generally, these findings illustrate how intrinsic mechanisms of excitability can be tuned by extracellular milieu, here via the dependence of *E*_*K*_ on [K^+^]_o_.

#### The role of *I*_*NaP*_ in modulating *I*_*CAN*_-based bistability

The persistent sodium current (*I*_*NaP*_) is well-known for amplifying neuronal excitability by providing a sustained depolarizing drive at subthreshold voltages. To explore its influence on motoneuron bistability, we examined its interactions with *I*_*CAN*_ using our computational model (Fig. 6). The two-parameter bifurcation map in Fig. 6B depicts the bistability range (color-coded as the hysteresis width *ΔI = I*_*up*_ *– I*_*down*_) as a function of *g*_*NaP*_ and *g*_*CAN*_. Overall, the bistable interval expands with increasing *g*_*NaP*_, indicating that *I*_*NaP*_ enhances the robustness of bistability. For instance, at moderately low *g*_*CAN*_ values (e.g., 0.9 mS/cm^2^; Fig. 6B), where *I*_*CAN*_ alone fails to produce bistability (Fig. 6C), elevating *g*_*NaP*_ from zero to 0.45 mS/cm^2^ uncovers a clear hysteresis interval (Figs. 6B, D-F), enabling the coexistence of silent and spiking states over a range of injected currents.

**Figure 6.**
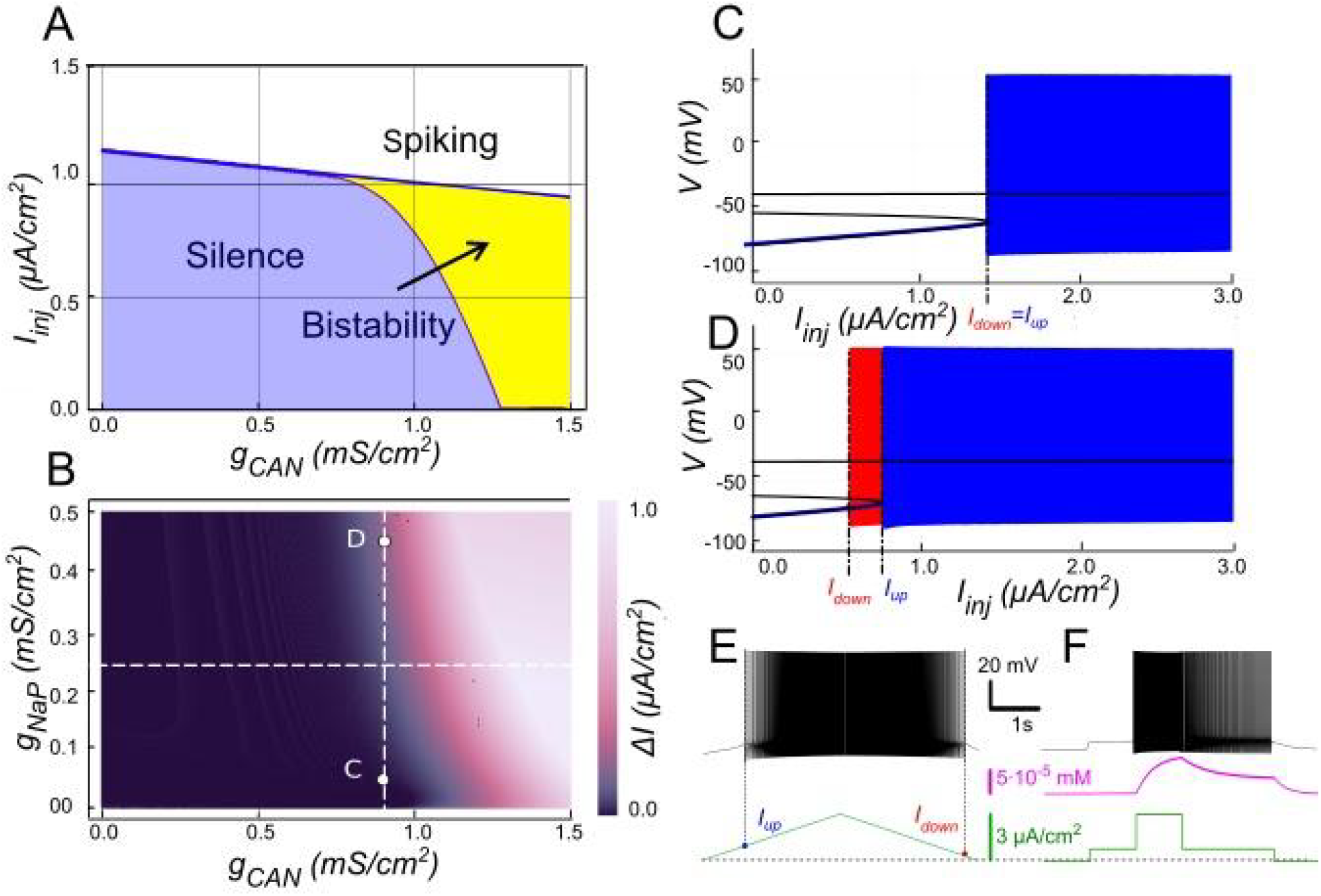
Modulation of *I*_*CAN*_-dependent bistability by *I*_*NaP*_. **A.**Bifurcation diagram similar to Fig. 3A, constructed for *g*_*KCa*_ = 0.5 *mS/cm*^*2*^, but at *g*_*NaP*_ = 0.25 *mS/cm*^*2*^ instead of zero. Note, that compared to Fig. 3A, bistability emerges at lower *g*_*CAN*_. **B**. Color-coded bistability range (*I*_*up*_ - *I*_*down*_) depending on *g*_*CAN*_ and *g*_*NaP*_ with *g*_*KCa*_ fixed at 0.5 *mS/cm*^*2*^. Black area corresponds to no bistability. *g*_*CAN*_ bifurcation value reduces as *g*_*NaP*_ increases, therefore an increase in *g*_*NaP*_ can lead to bistability emergence, as shown in **C** and **D**. If *g*_*CAN*_ = 0.9 *mS/cm*^*2*^, at *g*_*NaP*_ = 0 no bistability exists (**C**), but if *g*_*NaP*_ is raised to 0.45 *mS/cm*^*2*^, bistability emerges (**D**), as illustrated by ramp (**E**) and step current protocols (**F**).

This effect aligns with the core *I*_*CAN*_-driven positive feedback loop underpinning bistability: (i) spiking activity opens voltage-gated calcium channels, leading to Ca^2+^ influx; (ii) this influx triggers CICR, amplifying the cytosolic Ca^2+^ signal; (iii) elevated [Ca^2+^]_i_ then activates *I*_*CAN*_, and (iv) the resulting depolarisation accelerates spiking promoting additional Ca^2+^ entry and closing the self-reinforcing loop. Amplifying any element of this loop can elevate its overall gain, tipping the system toward bistability.

Here, *I*_*NaP*_ contributes by delivering a tonic depolarizing current that lowers the voltage threshold for spike initiation and Ca^2+^ entry. This facilitates the recruitment of *I*_*CAN*_ during the onset of activity and bolsters its ability to sustain the high-activity state once engaged, even at lower *g*_*CAN*_ levels. For example, at *g*_*CAN*_ = 0.9 mS/cm^2^ and *g*_*NaP*_ = 0.45 mS/cm^2^, the model displays robust bistability (Figs. 6D-F), whereas reducing *g*_*NaP*_ to 0 eliminates it (Fig. 6C). However, at lower *I*_*CAN*_ expression (e.g., at *g*_*CAN*_ = 0.5 mS/cm^2^), *I*_*NaP*_-evoked depolarization is insufficient to achieve the high-gain regime needed for bistability at any *g*_*NaP*_.

We tested these predictions pharmacologically in patch-clamp recorded lumbar motoneurons using veratridine, which at low micromolar concentrations enhances *I*_*NaP*_. Following a short depolarizing pulse, veratridine increased the amplitude and area of the sADP, while shortening its duration (Fig. 7A-D), indicating a stronger depolarizing tail when *I*_*NaP*_ is enhanced. Veratridine also facilitated bistability: self-sustained spiking activity were triggered from more hyperpolarized holding potentials, with *Vh* shifting from -56.9 mV to -61.6 mV (p < 0.01; Wilcoxon matched pairs test). Both the voltage range (*ΔV*) and the current range (*ΔI*) supporting bistability increased (Fig. 7E-H).

**Figure 7.**
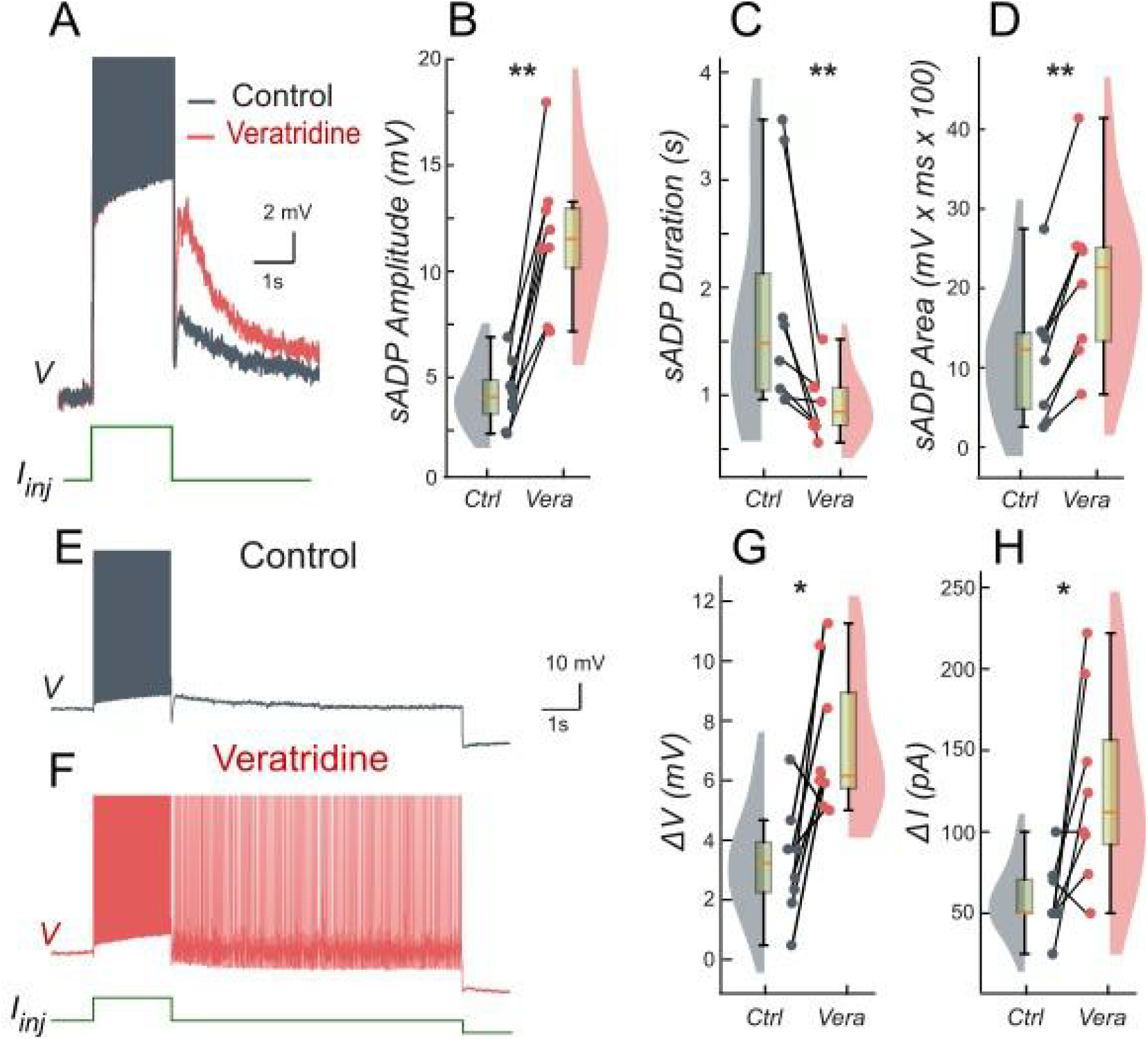
Persistent Na^+^ current (*I*_*NaP*_) facilitates the slow afterdepolarization and membrane bistability in lumbar motoneurons. **A**: Superimposed voltage traces recorded in the same motoneuron during a brief (2 s) depolarizing current step (bottom) under control conditions (black) and after bath application of veratridine (red). **B-D**: Quantification of the sADP amplitude (**B**), duration (**C**) and area (**D**). **E, F**: Voltage traces in response to a 2-s depolarizing pulse before (**E**) and after veratridine (**F**). **G, H:** Quantification of bistability through ΔV and ΔI. ΔV and ΔI represent the range of holding potentials and holding currents, respectively, over which self-sustained firing can be observed. Each is defined as the difference between the most depolarized and the most hyperpolarized value (potential for ΔV, current for ΔI) at which self-sustained firing is triggered or maintained (see Methods). Paired data from individual motoneurons (n = 8) are linked and overlaid on violin and box-and-whisker plots. *P < 0.05, **P < 0.01, two-tailed Wilcoxon signed-rank test.

Taken together, these findings support the model prediction that increasing *I*_*NaP*_ facilitates and broadens the operating window of *I*_*CAN*_-driven bistability by enhancing depolarization while *I*_*CAN*_ remains the principal maintenance mechanism once engaged. This synergy between *I*_*CAN*_ and *I*_*NaP*_ helps explain how modest changes in persistent Na? conductance can markedly reshape motoneuron firing regimes.

### Bistability based on *I*_*NaP*_ and the role of [K^+^]_o_

*I*_*NaP*_ could theoretically sustain bistability independently of *I*_*CAN*_ under conditions that maintain its activation between action potentials. For instance, if the inter-spike membrane potential remains above *I*_*NaP*_’s deactivation threshold, *I*_*NaP*_ would provide a continuous depolarizing drive, creating a self-reinforcing loop where subthreshold depolarization promotes spiking, and the resulting activity further engages *I*_*NaP*_ without full reset. This mechanism might enable the neuron to toggle between quiescent and self-sustained firing states purely through sodium-based persistence, highlighting a potential alternative pathway for bistability in scenarios where calcium-dependent processes are minimized or absent.

In our model, when *g*_*CAN*_ = 0, *I*_*NaP*_ alone could not support bistability in the parameter range examined (Fig. 8A). The bifurcation diagram (Fig. 8D) elucidates this. At low injected current (*I*_*inj*_ < 0.85 *μ*A/cm^2^), the system’s sole stable state is a low-voltage resting state (stable node). As *I*_*inj*_ surpasses this threshold, the node merges with a saddle point and subsequently ceases, leading to a stable limit cycle, which represents a spiking regime. A strong hyperpolarization follows each spike, dropping below the resting potential. This large post-spike hyperpolarization fully deactivates *I*_*NaP*_, explaining its inability to maintain bistability independently. Consequently, when the injected current is decreased, the silent regime reappears via the same bifurcation, demonstrating an absence of hysteresis.

**Figure 8.**
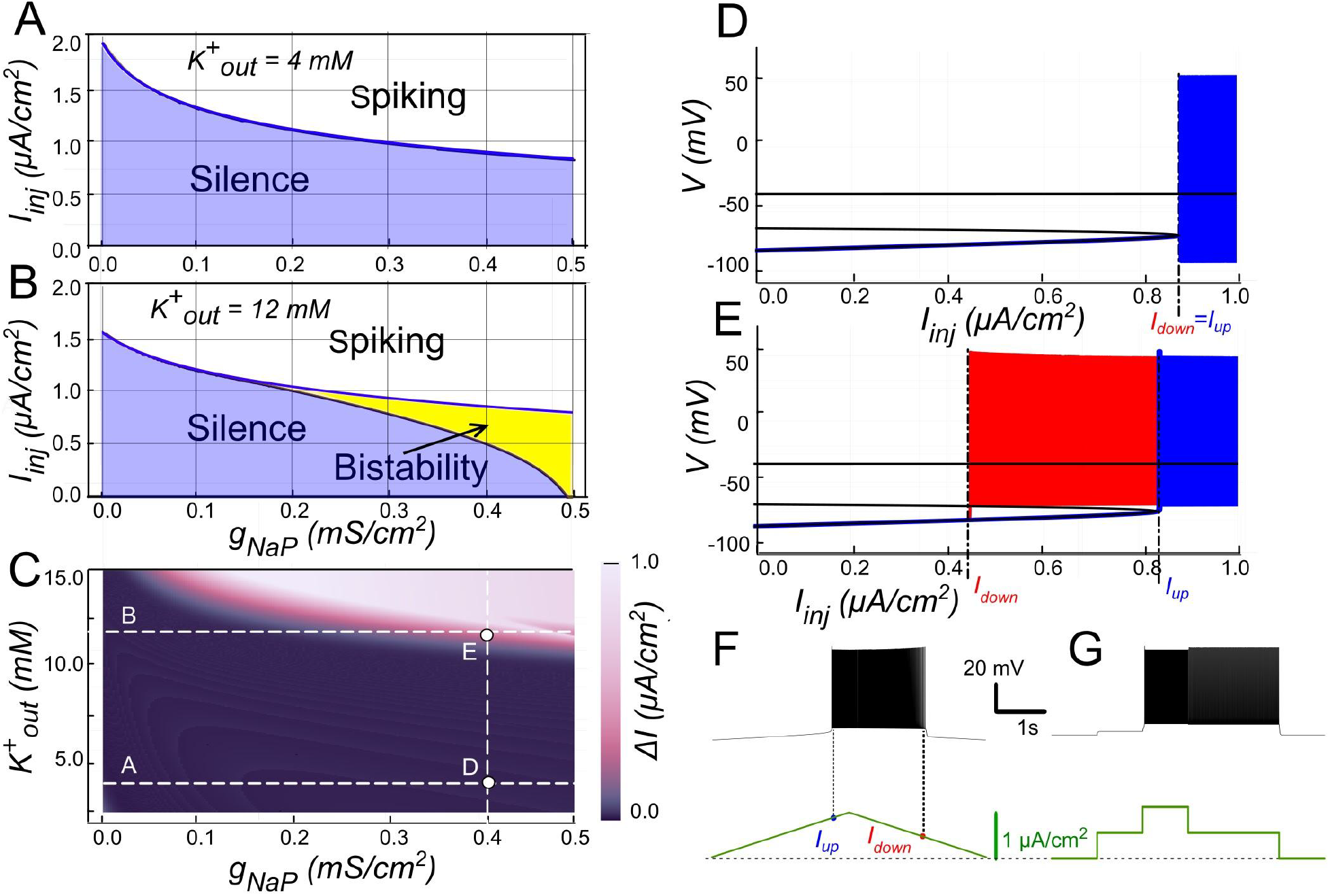
Bistability based on *I*_*NaP*_ and the role of [K^+^]_o_. **A.**Activity regimes of the model neuron depending on the injected current (*I*_*inj*_) and the conductance of persistent sodium current (*g*_*NaP*_) at baseline *K*^*+*^ extracellular concentration ([K^+^]_o_ = 4mM). At all values of *g*_*NaP*_ as the injected current changes, the model transitions from silence to spiking and back with no hysteresis which indicates no bistability. **B**. Activity patterns of the model neuron depending on the injected current and *g*_*NaP*_ conductance at [K^+^]_o_ = 12 mM. Once *g*_*NaP*_ exceeds approximately 0.15 mS/cm^2^, bistability emerges. **C**. Adjusting the sodium persistent inward current (*I*_*NaP*_) conductance (*g*_*NaP*_) and extracellular potassium concentration ([K^+^]_o_) in a model neuron reveals bistable regimes. The range of injected current where bistability occurs is shown in color, with black indicating no bistability. Higher [K^+^]_o_ levels require smaller *g*_*NaP*_ for bistability, suggesting that increased [K^+^]_o_ can induce bistability in neurons with otherwise insufficient *g*_*NaP*_ expression. **D**. at *g*_*NaP*_ = 0.4 mS/cm^2^ and [K^+^]_o_ = 4 mM no bistability is observed. **E**. However, as [K^+^]_o_ is increased to 12 mM, bistability emerges, as illustrated by ramp (**F**) and step current protocols (**G**).

The post-spike hyperpolarization is mediated by K^+^ outward currents and thus can be modulated by [K^+^]_o_, which sets the reversal potential *E*_*K*_. Raising [K^+^]_o_ depolarizes *E*_*K*_, thereby reducing the driving force of outward currents and diminishing the post-spike hyperpolarization. Under these conditions, *I*_*NaP*_ is less completely deactivated, allowing bistability to emerge (Figs. 8B, E).

As shown in Fig. 8E, when [K^+^]_o_ = 12 mM and *I*_*inj*_ crosses a value of 0.8 *μ*A/cm^2^, the resting state is lost through the same saddle-node bifurcation, but unlike at normal [K^+^]_o_, the trajectory joins a pre-existing spiking limit cycle with attenuated post-spike hyperpolarization (Figs. 8E, F). When *I*_*inj*_ is reduced, spiking persists down to 0.45 *μ*A/cm^2^, where the limit cycle intersects with the saddle point and disappears through a saddle-loop (homoclinic) bifurcation, and the system returns to resting state (Figs. 8E, F). Thus when *I*_*inj*_ falls between 0.45 and 0.85 *μ*A/cm^2^ both rest and spiking coexist, indicating a bistable regime (Fig. 8G). A distinctive feature of this type of bistability is that the resting potential lies below the voltage range of the limit cycle (spiking).

The presence and extent of bistable behavior depends jointly on *g*_*NaP*_ and [K^+^]_o_. At [K^+^]_o_ = 12 mM, bistability emerges once *g*_*NaP*_ exceeds ∼0.2 mS/cm^2^ (Fig. 8B), with the bistable interval widening as *g*_*NaP*_ increases. Mapping across parameters (Fig. 8C) shows that for *g*_*NaP*_ < 0.5 mS/cm^2^ bistability requires [K^+^]_o_ > 10 mM; conversely, the bistability *g*_*NaP*_ threshold decreases as [K^+^]_o_ increases. For instance, with *g*_*NaP*_ = 0.25 mS/cm^2^, bistability emerges once [K^+^]_o_ exceeds ∼12 mM.

Together, these results indicate that while under normal conditions *I*_*NaP*_ alone is insufficient to produce bistability, elevated [K^+^]_o_ reduces outward current-mediated hyperpolarization, potentially enabling bistable firing.

### The role of slowly inactivating potassium current (*I*_*Kv1*.*2*_)

Recent work indicates that the potassium current mediated by Kv1.2 channels (*I*_*Kv1*.*2*_) is prevalent in bistable motoneurons (Harris-Warrick et al., 2024), although its direct contribution to bistable behaviour has not been fully clarified. Kv1.2 channels inactivate very slowly during repetitive firing. In principle, such slow inactivation generates a positive feedback loop: as *I*_*Kv1*.*2*_ gradually decreases during ongoing spiking, the cell becomes more excitable and can persist in an active state. On the other hand, when the neuron is silent, Kv1.2 channels are fully open and *I*_*Kv1*.*2*_ can strongly oppose the initial depolarization.

To clarify *I*_*Kv1*.*2*_ contribution, we simulated our computational model while varying both *I*_*inj*_ and *I*_*Kv1*.*2*_’s maximal conductance, *g*_*Kv1*.*2*_. In these simulations, *I*_*Kv1*.*2*_ alone did not produce bistability between resting and spiking states. Instead, the model generated a regime of periodic bursting, alternating between spiking and quiescent phases, over an extremely narrow range of *I*_*inj*_.

A signature of *I*_*Kv1*.*2*_ is its effect on firing dynamics during constant depolarization: rather than stabilizing tonic firing, it induced a delayed excitation with ramping spike frequency (Bos et al., 2018). To isolate this effect, we removed both *I*_*CAN*_ and *I*_*NaP*_ (*g*_*NaP*_ = *g*_*CAN*_ = 0). Under these conditions (Fig. 9B), a rectangular current pulse elicited an initial fast depolarization followed by a slow secondary depolarization. If the stimulus was strong enough, this slow drift brought the neuron to a spiking threshold and then spiking began, with the firing rate progressively accelerating. This behavior reflects very slow inactivation kinetics of *I*_*Kv1*.*2*_. Initially, *I*_*Kv1*.*2*_ activates quickly, temporarily opposing depolarization, but then inactivates with a time constant of ∼2.5 s (see Methods), progressively reducing its own inhibitory effect and permitting further depolarization and firing.

**Figure 9.**
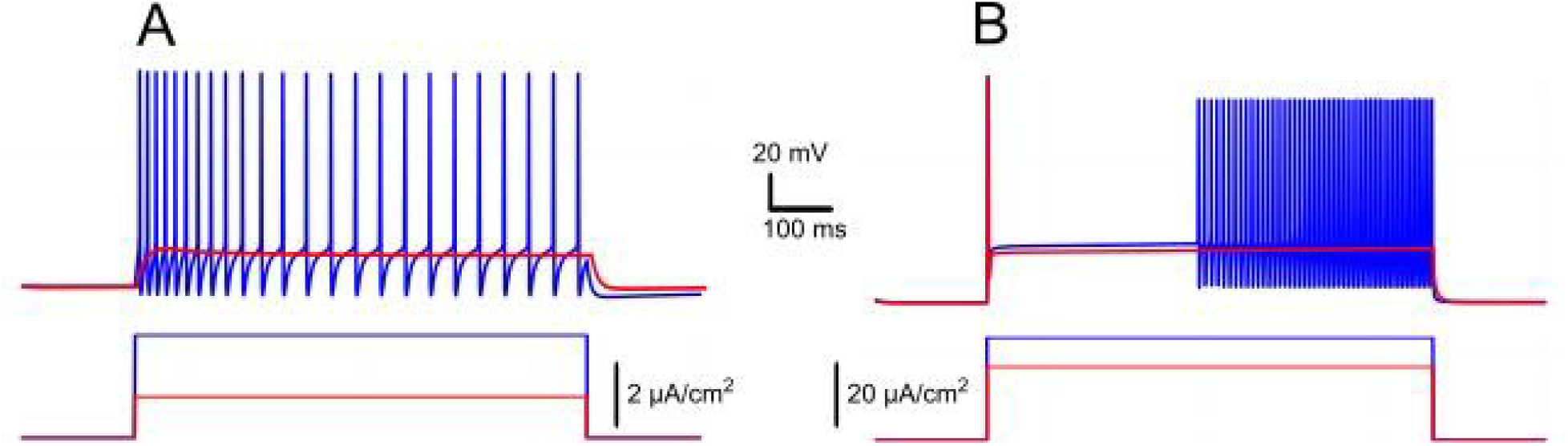
Ramp/delayed excitation versus spike frequency adaptation in response to rectangular current injection. **A.**Model response with *I*_*KCa*_ alone (*g*_*Kv1*.*2*_ = 0, *g*_*KCa*_ = 1 *mS/cm*^*2*^, *g*_*NaP*_ = *g*_*CAN*_ = 0), displaying spike frequency adaptation as *I*_*KCa*_ activation hyperpolarizes the neuron, reducing firing rate over time. **B**. Model response with *I*_*Kv1*.*2*_ alone (*g*_*Kv1*.*2*_ = 2 *mS/cm*^*2*^, *g*_*KCa*_ = 0, *g*_*NaP*_ = *g*_*CAN*_ = 0), showing delayed excitation and a ramping firing rate due to slow inactivation of *I*_*Kv1*.*2*_. The membrane potential exhibits an initial jump followed by gradual depolarization, leading to progressively increasing spike frequency.

By contrast, in absence of *I*_*Kv1*.*2*_ but with *I*_*KCa*_ present, the model showed spike frequency adaptation (Fig. 9A). Here, firing decreased over time because *I*_*KCa*_ hyperpolarized the membrane. When both currents (*I*_*Kv1*.*2*_ and *I*_*KCa*_) were present, the firing dynamics (whether it shows ramping or adaptation) depended on their relative balance. This interaction can lead to complex dynamics where the slow inactivation of *I*_*Kv1*.*2*_ competes with the calcium-dependent potassium currents tendency to stabilize or reduce firing frequency.

## Discussion

In this study, we used a single-compartment computational model of spinal motoneurons to dissect the ionic mechanisms underlying bistability. Our results identify a minimal core mechanism based on the synergistic interactions among *I*_*CaL*_, CICR, and *I*_*CAN*_ strongly modulated by *I*_*KCa*_, *I*_*NaP*_ and [K^+^]_o_. These findings offer new insights into how bistable firing is generated and regulated in motoneurons.

### Motoneuron bistability critically depends on the *I*_*CaL*_-CICR-*I*_*CAN*_ loop

Early studies attributed plateau potentials and bistability to persistent L-type calcium currents (Schwindt & Crill, 1980; Hounsgaard, Hultborn, et al., 1988; Hounsgaard, Kiehn, et al., 1988; Hounsgaard & Mintz, 1988; Hounsgaard & Kiehn, 1989; Hounsgaard & Kjaerulff, 1992; Svirskis & Hounsgaard, 1997), a view reinforced by computational models requiring dendritic *I*_*CaL*_ to replicate these bistable behaviors (Booth & Rinzel, 1995; Booth et al., 1997; Carlin et al., 2000; ElBasiouny et al., 2005; Bui et al., 2006; Carlin et al., 2009; Kim & Jones, 2011). Immunohistochemical data further supported this interpretation by revealing the dendritic distribution of L-type Cav1.3 channels (Simon et al., 2003; Zhang et al., 2008). More recent findings have revealed a complementary mechanism, in which *I*_*CaL*_ primarily acts as a trigger, while other currents, notably the *I*_*CAN*_, play a central role in mediating plateau potentials and bistability (Bos et al., 2021), as observed in motoneurons from neonatal and young adult murine models (Harris-Warrick et al., 2024).

Our computational findings reinforce and extend this view. The model shows that initial Ca^2+^ entry through *I*_*CaL*_ is insufficient to sustain bistability due to fast Ca^2+^ removal from the cytosol. The necessary amplification arises from CICR, linking *I*_*CaL*_-mediated Ca^2+^ influx to intracellular stores. Accordingly, CICR surpasses the capacity of Ca^2+^ pumps to rapidly clear cytoplasmic Ca^2+^, and therefore enables sustained intracellular Ca^2+^, long-lasting *I*_*CAN*_ activation and bistability. Consistent with this, motoneuron bistability is abolished when CICR is inhibited, even though *I*_*CaL*_ currents are still present (Bouhadfane et al., 2013; Bos et al., 2021; Harris-Warrick et al., 2024). Together, *I*_*CaL*_, CICR, and *I*_*CAN*_ emerge from the findings as a functional triad instrumental in motoneuron bistability. As such, the triad may provide a general feed-forward mechanism for plateau generation across different structures of the CNS, owing to the fact that *I*_*CAN*_ also supports motor, sensory and memory-related plateaus (Fraser & MacVicar, 1996; Morisset & Nagy, 1999; Di Prisco et al., 2000; Yan et al., 2009; Toporikova & Butera, 2011; Jasinski et al., 2013).

### Suppression of *I*_*CAN*_ -based bistability by *I*_*KCa*_

The model adds a key regulatory layer to motoneuron bistability by clarifying the interplay between *I*_*CAN*_ and *I*_*KCa*_. *I*_*KCa*_ dominance inhibits *I*_*CAN*_ -driven plateaus and prevents their formation. In line with the model, reducing *I*_*KCa*_ with apamin facilitates the expression of bistability and can unmask latent plateau potentials (Hounsgaard & Mintz, 1988; Hounsgaard & Kiehn, 1993).The relative contributions of *I*_*CAN*_ and *I*_*KCa*_ may tune the propensity to bistability across motoneuron subtypes. Bistability appears more frequently observed in large motoneurons (Harris-Warrick et al., 2024), a finding that, according to the assumption, would reflect a higher *I*_*CAN*_ /*I*_*KCa*_ ratio. This interpretation aligns with established physiological distinctions where large motoneurons, in line with size-dependent differential expression of SK channels (Deardorff et al., 2013), display brief AHPs, whereas small motoneurons exhibit prolonged AHPs (Kernell, 1965; Gustafsson & Pinter, 1984). In addition, large motoneurons generate stronger *I*_*CAN*_ than small motoneurons (Harris-Warrick et al., 2024).

Our model also identifies [K^+^]_o_ as a critical factor influencing *I*_*KCa*_ efficacy. Elevating [K^+^]_o_ depolarizes *E*_*K*_, weakening *I*_*KCa*_ and shifting the balance towards *I*_*CAN*_, thereby enhancing bistability (Fig. 4). Disruption of K^+^ buffering through astrocytic Kir4.1 dysfunction in spinal cord injury (Olsen et al., 2010; Benson et al., 2023; Barbay et al., 2024), is likely to elevate [K^+^]_o_, paralleling findings from epilepsy studies (Djukic et al., 2007; Tong et al., 2014). A subsequent shift towards *I*_*CAN*_ is predicted to strengthen bistability and worsen spasticity (Bennett et al., 2001; C. Brocard et al., 2016).

### Facilitation of *I*_*CAN*_ -mediated bistability by *I*_*NaP*_

While *I*_*CAN*_ is the main driver of bistability, *I*_*NaP*_ acts as an essential modulator, extending the conditions under which bistability occurs. In our simulations, increasing *I*_*NaP*_ lowers the threshold for *I*_*CAN*_-mediated bistability, enabling sustained firing even when *I*_*CAN*_ alone is insufficient (Fig. 6). Due to subthreshold depolarization (Crill, 1996) that enables repetitive spiking (Kuo et al., 2006), *I*_*NaP*_ biases the system towards *I*_*CaL*_-CICR-*I*_*CAN*_ engagement and plateau generation. In line with this role, riluzole, an established inhibitor of *I*_*NaP*_, reliably suppresses self-sustained firing in bistable motoneurons (Bouhadfane et al., 2013; Drouillas et al., 2023). At first glance, this might imply that plateau potentials depend directly on *I*_*NaP*_. However, the persistence of TTX-resistant sADP after rizulole application (Bouhadfane et al., 2013; Drouillas et al., 2023) suggests that *I*_*CAN*_ provides the essential substrate for bistable behavior, while *I*_*NaP*_ serves as a facilitator. The *I*_*NaP*_ and *I*_*CAN*_ interaction extends beyond motoneurons; in the preBotzinger complex, for example, the two currents cooperate to produce rhythmic bursting (Jasinski et al., 2013; Phillips et al., 2019, 2022).

Under physiological conditions, *I*_*NaP*_ alone unlikely supports bistability because hyperpolarizing potassium currents deactivate *I*_*NaP*_ between spikes (Fig. 8). Elevating [K^+^]_o_ mitigates this hyperpolarization by depolarizing *E*_*K*_, and allows *I*_*NaP*_ to create bistability at higher *g*_*NaP*_ (Fig. 8C). This has been clearly demonstrated in our simulations, where a pure *I*_*NaP*_ -based bistability emerges as [K^+^]_o_ approaches or exceeds 12 mM. This scenario may be relevant in SCI, where *I*_*NaP*_ is enhanced (Bennett et al., 2001; Li et al., 2004; Harvey et al., 2006; C. Brocard et al., 2016). In conjunction with high [K^+^]_o_, this enhancement can strengthen bistability and promote spasticity. Consequently, riluzole (Rilutek©), originally developed for ALS, is being explored to target spasticity in SCI patients (Cotinat et al., 2023).

### The role of slowly inactivating potassium current (*I*_*Kv1*.*2*_)

In spinal motoneurons, *I*_*Kv1*.*2*_ imposes an initial brake on excitability and then relaxes over seconds, yielding delayed spiking and a characteristic ramping of discharge during sustained depolarization (Bos et al., 2018). Because this property scales with motoneuron size, larger α-motoneurons show stronger delayed excitation and ramping (Harris-Warrick et al., 2024). Yet, despite the greater prevalence of bistability in larger motoneurons, evidence for a generative role of Kv1.2 remains elusive. Our simulations clarify this point. Varying *I*_*Kv1*.*2*_ in isolation never produced robust switching between silent and self-sustained firing states. Instead, it generated narrow-band bursting around a tight input window, while the hallmark hysteresis of bistability was absent. Mechanistically, *I*_*Kv1*.*2*_ lacks the positive feedback needed to maintain a depolarized up-state. It is an outward conductance that weakens with use, modulating access to the plateau regime but not providing the sustaining inward drive. In contrast, stable bistability in both our model and experiments requires the *I*_*CaL*_-CICR-*I*_*CAN*_ triad, a view reinforced by the identification of Trpm5 as the principal Na^+^-permeable carrier of *I*_*CAN*_ underlying motoneuron plateaus (Bos et al., 2021). When TRPM5/*I*_*CAN*_ is suppressed, slow afterdepolarization and plateaus collapse even though *I*_*Kv1*.*2*_ is intact, directly demonstrating that Kv1.2 is permissive rather than generative for bistability.

### Serotonin and bistability

Brainstem-derived monoamines are central to motoneuron excitability and bistability (Heckman et al., 2008). In decerebrate cats, bistability depends on descending monoaminergic drive (Conway et al., 1988; Hounsgaard, Hultborn, et al., 1988; Lee & Heckman, 1998b, 1999). Acute spinalization removes these inputs and thereby reduces bistable properties of motoneurons, whereas monoamine reintroduction restores plateau potentials (Hounsgaard, Hultborn, et al., 1988). Among these modulators, serotonin is especially effective. In vertebrates, exogenous serotonin enhances excitability and bistability (Hounsgaard & Kiehn, 1985, 1989).

Mechanistically, serotonin acts primarily through 5-HT_2_ receptors to amplify ionic currents that promote bistability. By increasing dendritic L-type calcium currents (Hounsgaard & Kiehn, 1989; Perrier & Hounsgaard, 2003; Perrier & Delgado-Lezama, 2005; Perrier & Cotel, 2008) it can promote calcium build-up and *I*_*CAN*_ activation as described in our model. Serotonin also shifts *I*_*NaP*_ activation toward hyperpolarized potentials, thereby amplifying neuronal excitability (Li & Bennett, 2003; Harvey et al., 2006). The shift enhances *I*_*NaP*_ and thus facilitates *I*_*CAN*_-mediated bistability. Finally, serotonin also reduces outward currents, notably *I*_*KCa*_, thereby facilitating high-frequency firing (Grunnet et al., 2004). This serotonin-evoked reduction of *I*_*KCa*_ may lead to bistability in spinal motoneurons (Hounsgaard, Kiehn, et al., 1988; Hounsgaard & Kiehn, 1993), as demonstrated in our present modeling and experimental results.

The role of serotonin becomes especially evident during SCI, not because of its direct action, but rather because of its sudden loss following disruption of descending inputs. Initial serotonin depletion produces motoneurons hypofunction, otherwise known as “spinal shock” (Schadt & Barnes, 1980), mirroring the loss of bistability in our model upon blockade of *I*_*CAN*_ or CICR. However, with time, excitability and plateau potentials re-emerge, representing an electrophysiological correlate of chronic spasticity and hyperreflexia (Bennett et al., 2001). The rebound is attributed to plasticity in serotonin receptor signaling. Most notably, 5-HT_2B_ and 5-HT2_C_ receptors become constitutively active which restores persistent inward currents (e.g., *I*_*NaP*_) and plateau firing even without serotonergic input (Li & Bennett, 2003; Murray et al., 2010, 2011; D’Amico et al., 2013; Tysseling et al., 2017). Consistent with this, our model shows that increased *I*_*NaP*_ strengthens *I*_*CAN*_-driven bistability.

## Limitations and future research directions

Dendritic calcium currents have been repeatedly linked to motoneuron plateaus and bistability (Booth et al., 1997; Carlin et al., 2000; Simon et al., 2003; ElBasiouny et al., 2005; Bui et al., 2006; Zhang et al., 2008). Our single-compartment computational model reproduces key features of motoneuron bistability, yet its simplified architecture cannot capture the full spatial complexity of motoneurons. By collapsing dendrites into a one compartment, our results indicate that a single-compartment approximation can support bistability (Booth et al., 1997; Carlin et al., 2000; ElBasiouny et al., 2005; Bui et al., 2006). They do not, however, exclude the possible specific role of dendrites. Indeed, dendritic *I*_*CaL*_ might amplify calcium signaling and could shape the expression and robustness of plateaus.

Our future work will include the construction and use of multi-compartment models to explore the role of spatial distribution of *I*_*CaL*_, CICR, and *I*_*CAN*_ for testing how dendrite-versus soma-localized mechanisms affect bistability. In this context, establishing the precise subcellular localization of *I*_*CAN*_ channels remains a priority but progress is limited by the lack of highly specific antibodies to TRPM5, the presumed molecular correlate of *I*_*CAN*_ (Bos et al., 2021). Our analysis centers at L-type calcium channels only, but motoneurons also express T-type and N-type calcium channels (Umemiya & Berger, 1995; Hivert et al., 1995; Viana et al., 1997). Since these currents can complement or replace L-type currents in generating plateaus (Bouhadfane et al., 2013), future models may be instrumental in evaluating their contributions.

In summary, our minimal model captures the core features of bistable firing, with the ICaL-CICR-ICAN triad emerging as a central mechanism, while leaving open the contribution of dendritic processes. Future anatomically detailed models and experiments will be needed to resolve the effects of channels’ spatial distributions across motoneuron compartments.

## Acknowledgements

This work was supported by the ANR MotoBis grant (ANR-24-CE16-1548-01) to F.B. and the NIH/NINDS grant (R01 NS130799) to I.A.R.

